# Effects of an increase in water temperature on inter- and transgenerational plasticity reveal a short-term metabolic and phenotypic memory in an aquatic plant species

**DOI:** 10.64898/2026.07.06.736556

**Authors:** Grégoire Loupit, Mélissa Sancharme, Pierre Petriacq, Josep Valls Fonayet, Anne-Kristel Bittebiere

## Abstract

Transgenerational plasticity can shape plant phenotype and influence plant response to environmental changes in interaction with the current conditions. While how past stress interact with either current optimal or stress conditions is increasingly documented within a single plant, transgenerational plasticity remains particularly poorly understood especially at the metabolome level. In our study, we investigated whether heat stress induces transgenerational metabolic and phenotypic modifications along two successive clonal ramet generations of the sub-Antarctic aquatic plant *Limosella australis*. We performed untargeted metabolomics approaches and measured morphologic and performance traits, to assess both transgenerational plasticity of the metabolome and the phenotype. We found that heat stress remodelled the metabolic profile and influenced the foraging strategy of our clonal plant, and that some of these metabolic changes persisted into the first clonal generation. This one therefore adopted an intermediate growth strategy, even though culture conditions were optimal. By comparing differentially accumulated features between daughter ramets from heat stressed mother ramets and from unstressed mother ramets, we identified common and specific metabolites accumulation to heat stress response, belonging to diverse compound families. However, we did not observe any adaptative advantage and any metabolic imprint during another heat stress applied on the second clonal generation. This work provides especially new clues into how plant metabolome integrates and transfers previous stressed clonal generation’s information.

## Introduction

As sessile organisms, plants have to deal with stress through their plasticity—allowing a genotype to express different phenotypes under different environmental conditions (Pigliucci, 2001). However, genotype × environment interaction does not fully determine the expressed phenotype, as plant past experiences also influence its responses, indicating a plant memory (Trewavas, 2003; Conrath *et al*., 2006; Bruce *et al*., 2007; Auge *et al*., 2023). Indeed, plant phenotype can be modulated by individual (somatic memory) and parental (intergenerational and transgenerational memory) experiences, by storing molecular and chemical information (Herman & Sultan, 2011; Paszkowski & Grossniklaus, 2011; Lämke & Bäurle, 2017; Sharma *et al*., 2022; Auge *et al*., 2023; Pissolato *et al*., 2024). So far, somatic memory is considered as short-term (it does not persist in the next generation), transmitted only by mitotic cell division (Sharma *et al*., 2022), while intergenerational memory (transmitted from parents to offspring) and transgenerational memory (at least transmitted from grandparents to offspring) are related to long-term, transmitted through meiosis and sexual reproduction (Slaughter *et al*., 2012). In clonal plants (*sensu* Van Groenendael *et al*., 1996), mother memory can however be transmitted in daughters, through mitosis, during clonal propagation (Münzbergová & Hadincová, 2017; González *et al*., 2018; Zhang *et al*., 2021), involving mechanisms common to somatic memory. As clonal plants dominate most plant communities (Klimeš & Klimešová, 1999), understanding the mechanisms underlying their plastic responses, in particular how these are influenced by past environmental conditions, is essential. Yet, studies dealing with inter- and transgenerational plasticity investigated phenological and morphological responses (González *et al*., 2016, 2018; Münzbergová & Hadincová, 2017; Quan *et al*., 2022) without considering the metabolic status of individuals, although it seems responsible for response trait expression (Hemme *et al*., 2014; Serrano *et al*., 2019).

In nature, stress triggers metabolome reprogramming of plant individuals; exposure to drought, heat, and cold or oxidative stress activates defence mechanisms, and can enhance individual future tolerance (Hilker & Schmülling, 2019; Sharma *et al*., 2022). More specifically, higher temperatures induce fluctuations in both primary and secondary metabolite concentrations (Fraire-Velázquez & Balderas-Hernández, 2013). Sugars or amino acids accumulate as osmoprotectants (Khan *et al*., 2020), and secondary metabolites, like phenolic compounds, participate as antioxidants (Agati *et al*., 2012). Many metabolites are also involved in heat stress signalling pathways and, together with stress hormones (e.g. ethylene or abscisic acid), modify molecular processes (Wahid *et al*., 2007). However, although there is likely common response mechanisms to heat stress across plant species, other metabolite accumulations can be plant species dependent (Mandal *et al*., 2012; Benina *et al*., 2013).

The individual memory of past stress can be initiated by this metabolome reprogramming. Although there is little evidence in the literature (but see Benina *et al*., 2013; An *et al*., 2013; Hemme *et al*., 2014; Wedeking *et al*., 2018; Serrano *et al*., 2019), metabolic changes can actually persist between two episodes of stress i.e., during the recovery period (Schwachtje *et al*., 2019). Several metabolites can indeed remain in higher content, stored as active or inactive compounds (Pastor *et al*., 2013), such as amino acids (Benina *et al*., 2013; An *et al*., 2013; Hemme *et al*., 2014; Wedeking *et al*., 2018; Serrano *et al*., 2019) and primary metabolites like TCA intermediates or other central compounds (Mandal *et al*., 2012; Benina *et al*., 2013; Hemme *et al*., 2014; Serrano *et al*., 2019; Sharma *et al*., 2019; Olas *et al*., 2021). These metabolites would then constitute potential candidates of somatic, inter- and transgenerational memories, and should be found in daughters and granddaughters of stressed mothers.

Somatic memory of plant metabolome induces, when another stress occurs, a more efficient**—**faster and/or more intense**—**individual response (Charng *et al*., 2023). As a consequence, in *Arabidopsis sp*. for example, individuals exposed to a first episode of heat stress, tolerated a second episode much better than non-exposed plants (Serrano *et al*., 2019). However, little is known on whether inter- and transgenerational memories of plant metabolome similarly promote daughter and granddaughter performance in clonal plants [but see Bhatt *et al*. (2026)]. In parallel, in the clonal eelgrass *Zostera marina*, heat stress increased shoot length of daughters of which mothers were previously exposed, but also reduced their relative growth rate (DuBois *et al*., 2020). Metabolome effects on plant performance may thus actually occur through the modification of plant functional traits, although this hypothesis remains to be thoroughly tested.

The aim of our study was to characterise inter- and transgenerational metabolic responses to heat stress in clonal plants, and the consequences for their growth, and morphological and performance traits. This controlled experiment focused on *Limosella australis*, a macrophyte species found in the freshwater ponds of the Iles Kerguelen (sub-Antarctic region), currently experiencing rapid and intense climate warming (Leihy *et al*., 2018; Douce *et al*., 2023). We exposed plants to either control (13 °C) or heat stress (23 °C) conditions, across three clonal generations (mother, daughter, and granddaughter), with or without a recovery period (Fig. 1). On daughters and granddaughters only (respectively R2 and R3 hereafter), we measured functional traits, speed of clonal propagation, and individual dry weights, and analysed their metabolic profile using an untargeted metabolomics approach (both in negative and positive ionisation). Through this study, we specifically tested the following hypotheses:

1. Heat stress (23°C) induces individual metabolism reprogramming and triggers both primary and secondary metabolism.
2. During recovery period, daughters maintain metabolic changes resulting from mother heat stress, suggesting the existence of an intergenerational memory.
3. Granddaughters of stress-exposed grandmothers differentially express metabolic changes in response to heat stress, consistently to a transgenerational memory.
4. Heat stress induces morphological and growth responses in mothers that can be transmitted in their daughters and granddaughters, enhancing their performance under a new stress episode, which highlights inter- and transgenerational plasticity through clonal generations.

**Figure 1:**
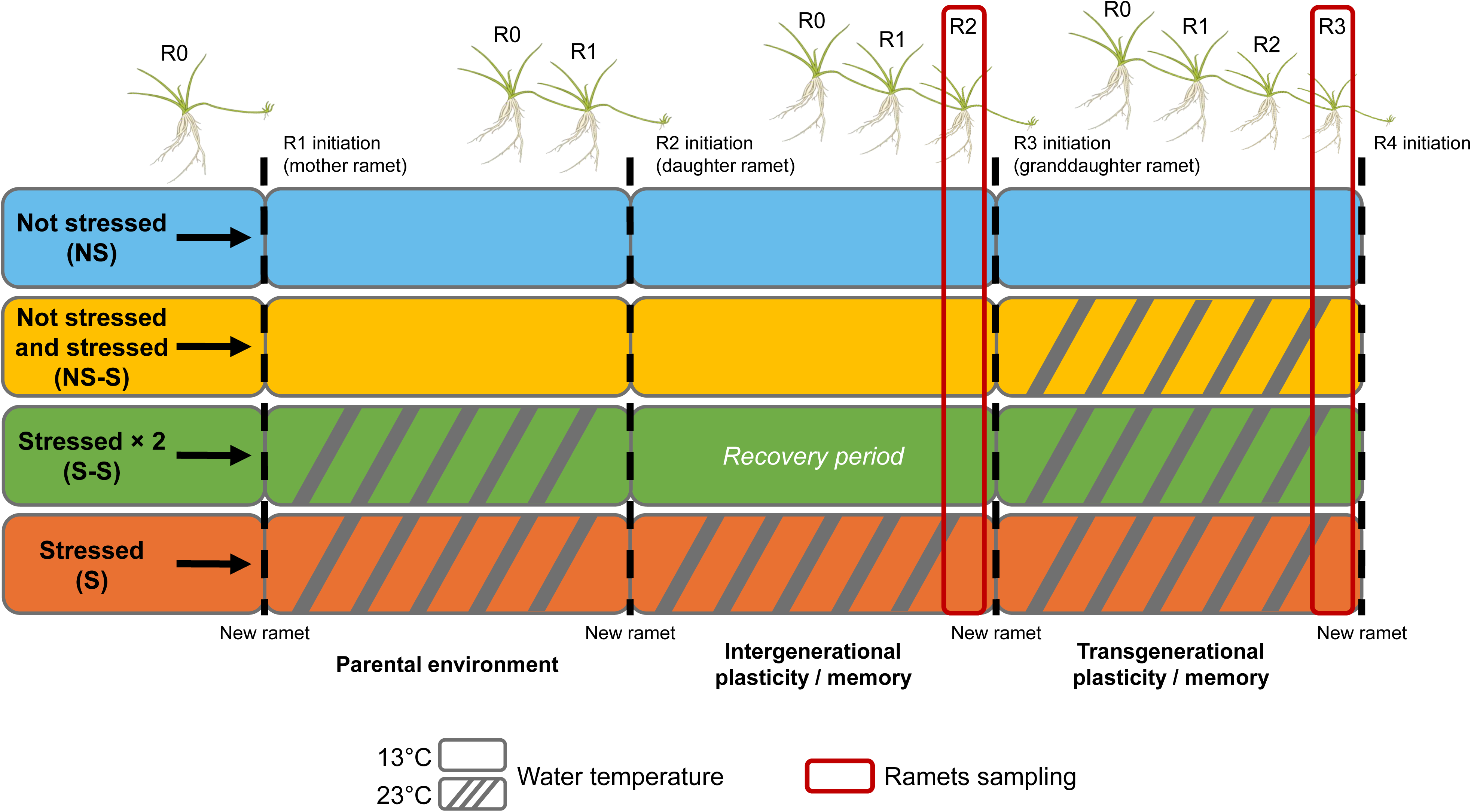
Experimental design showing the different growing conditions (water temperature undergone by each ramet generation for the four treatments). Each clone was moved to a new temperature condition when we observed the appearance of a new clonal ramet. Sampling, for morphologic traits, dry weight and metabolomics analysis, was done on daughter (R2) and granddaughter (R3) ramets when we observed the appearance of a new clonal ramet (R3 and R4 respectively).

## Materials and methods

### 1. The natural ecosystem of *Limosella australis*

Our controlled experiment was based on the species *Limosella australis* that live in freshwater ponds of the Iles Kerguelen (Southern Indian Ocean, 49°20’00’’S, 69°20’00’’E). These freshwater ponds accumulate nutrients from marine fauna (Smith, 2008) and are experiencing considerable warming because of climate change (Leihy *et al*., 2018). Since 1960, annual air temperatures have risen by 1.3°C, accompanied by a decrease in precipitation both in terms of quantity and frequency, and a reduction in frost days (Frenot *et al*., 2006). Aquatic plant species, i.e. macrophytes, from the Iles Kerguelen, adapted to low temperatures**—**typically around 13 °C during the day in summer in freshwater ponds (Douce *et al*., 2023)**—**are thus particularly vulnerable to the current warming, and may experience heat stress *in situ*. *L. australis* is a perennial and clonal species producing new daughter ramets (short erected shoot with its leaves and roots, forming rosettes) connected through plagiotropic stems forming a clonal network, allowing it to colonise freshwater ponds. We can then easily differentiate each generation of ramet.

### 2. Plant material

Seeds of *L. australis* were collected from freshwater ponds at different locations of the Iles Kerguelen to maximise genetic variability. These were germinated and new individuals were cultivated to clonally propagate them, in pots on a 50% sand / 50% compost (N = 250, P_2_O_5_ = 120, and K_2_O = 80 mg.L^−1^) substrate, submerged in trays filled with water. These cultures last for two years under controlled conditions (at 13 °C with 12/12 h of light/dark, at 3000 lux, and with air bubblers). This allowed us to minimise maternal effects and to produce numerous ramets of the same clone (i.e., of the same genotype). These are then assigned to different treatments in our experiment, to control for genotype effects on the studied response.

### 3. Study design

Our experiment was built (i) to characterise metabolic changes in daughters and in granddaughters resulting from inter- and transgenerational metabolic memory of stress-exposure in mothers; (ii) and to determine how it subsequently affects the growth, and morphological and performance traits on both daughter and granddaughter ramets.

To achieve these objectives, we selected 24 ramets (R0) of eight *L. australis* genotypes, and cultivated them in separate pots (8 cm width × 8 cm length × 7 cm height; 50% sand and 50% compost; N = 250, P_2_O_5_ = 120, and K_2_O = 80 mg.L^−1^; 12-hour light-dark cycles, 3000 lux and with air bubblers). These 24 ramets were selected to be of the same age and size and were maintained in the same conditions (13 °C, mean water temperature in freshwater ponds during summer days) until that new ramets (R1 hereafter) were produced. Just before these R1 began to develop (when the internode bud initiates a new ramet), we randomly assigned and transferred six R1 ramets of each genotype, in each of the four following treatments: ‘Not Stressed’ (NS, control), ‘Not Stressed and stressed’ (NS-S), ‘Stressed and stressed’ (S-S, including a recovery period between two episodes of heat stress) and ‘Stressed’ (S, no recovery period) (Fig. 1). Then, depending on the tested treatments, new clones (resulting from the clone propagation of R1) were transferred into different conditions of temperature, at each new ramet initiation. If several connections were produced from the same ramet, only one was preserved, and the others were manually cut. Half of the clones were harvested at the initiation of R3 to sample R2 adult ramets, and the other half at the initiation of R4 to sample R3 adult ramets.

### 4. Characterisation of growth, and measurements of morphological and performance traits

First, we counted the number of days needed to initiate ramets R3 from R2, and R4 from R3, to measure the speed of clonal propagation i.e., growth.

Second, on R2 and R3, we additionally measured morphological and performance traits. The maximum leaf length and the maximum root length were measured as the functional characteristics of above- and below-ground organs. Also, we measured the length of stolon internodes starting from R2 or R3, as descriptor of their clonal dispersal. Immediately after these morphological measurements, these ramets were frozen in liquid nitrogen. Samples were then stored at -80 °C for further analyses. In addition, we measured the ramet dry weights as an indicator of performance (including ramet leaves, roots, stem, and internode).

### 5. Metabolite extraction

Before the metabolomics analyses, the three ramets of the same generation (R2 or R3) and the same genotype that endured the same treatment, were pooled together to have enough plant matter .

Lyophilised samples were weighted and grounded to powder with two stainless steel beads in their own tube using a 2010 Geno/Grinder® (SPEX® SamplePrep) during 3 × 1 min (with 15 s of rest) at 29.2 Hz (1750 strokes/min). A methanol extraction was performed by adding 600 µL of MeOH in each tube, plonged in an ultrasound bath during 15 min and centrifuged at 10,000 g during 10 min at 4 °C. Finally, supernatants were recovered and dried using a Concentrator plus / Vacufuge® plus at 30 °C.

### 6. Untargeted metabolomics analyses

Dry extracts were recovered in 100 µL of MeOH. Then, the untargeted metabolomics analyses were performed with a Vanquish UHPLC (Thermo Scientific) coupled to an Orbitrap instrument (Qex+, Thermo Scientific). One microliter of sample was injected on a Phenomenex Luna® Omega Polar C18 column (50 × 2.1 mm, 1.6 µm) at 40 °C and a gradient of solvent A (milliQ water – 0.1 % formic acid) and solvent B (acetonitrile – 0.1% formic acid) with a flow of 0.5 mL.min^−1^ was used. The gradient elution was set as follows for solvent B: 0-7.5 min: 1-36%; 7.5-14 min: 36-80%; 14-15 min: 80-95%; 15-17 min: 95%; 17-18.5 min: 1%. The source conditions were as follows: Spray voltage: 3000 V; Sheath gas: 45 a.u; Auxiliary gas: 15 a.u; Capillary temperature: 320°C; Probe heater temperature: 250°C; S-lens RF level: 100. The experiments were done in full scan (mass range: 70-1050 m/z)—data depending on MS/MS. The MS data were acquired in negative polarity at 140.000 FWHM resolution with an automatic gain target set at 3e6 and maximum IT of 100 ms. The MS2 fragmentation data were acquired at 35.000 FWHM with top two precursors and normalised collision energies of 15, 30 and 40 using a dynamic exclusion of 3s.

Before injection, all samples were randomised and then analysed with 10 Quality Control (QC) and extraction blanks (QC samples were injected every 10 runs and were used to correct the potential signal drift). All samples were analysed two times, in negative and positive mode ionisation (Supplementary File 1). For each run of analysis (positive and negative mode), raw data were processed by using MS-DIAL v4.9 (Tsugawa *et al*., 2015) configured with different parameters (Supplementary File 2 and 3) with LOWESS normalisation method based on QC samples. Feature annotations were attributed based on MS1 spectra and MS2 DDA fragmentation information using the MetaboHUB AgroMix in-house chemical database (generously borrowed from Guillaume Marti, AgroMix MetaboHUB-MetaToul, France). Finally, we retained 3267 and 5942 features (after removing features with SN < 10 and coefficient of variation of QC samples > 30%), where 99 and 132 matched with MS1 and MS2, 1829 and 3436 suggested features matched with MS1 only, and 1339 and 2374 unknown (no match) in negative and positive mode respectively. ClassyFire was used to automatically produce chemical ontology through the InChiKeys of annotated features identified (Djoumbou Feunang *et al*., 2016).

Finally, samples were normalised by their weights, and QC and blanks were normalised by the mean weight of all pooled samples (Supplementary Data 1 and 2).

### 7. Statistical analyses

#### Trait data

To assess the effects of the different treatments on growth, and morphological, and performance traits of R2 or R3 ramets of *Limosella australis*, we used Linear Mixed-Effects Models (LMM - package *lme4*, version 1.1-36) (Bates *et al*., 2015). The treatment was used as fixed factor, and the clone genotype was used as random factor. This allowed us to assess trait plastic response as genotype reaction norm. For number of days to initiate a new clonal ramet, a square-root transformation was applied. For internode length only, we added the ramet biomass as covariate in the model (without interaction with treatment) to evidence a possible active response of the ramet instead of a passive response that could be related to an overall decrease in the ramet growth. Then, we performed ANOVA or ANCOVA (type II with Kenward–Roger degrees of freedom) based on the built models (Table 1) (package *stats*, version 4.4.1). We then calculated Estimated Marginal Means (Adj. means) and standard error from LMM using the *emmeans* package (version 1.10.4) (Lenth, 2017), correcting random factor and covariate effects. When the effect of the treatment was significant, we performed Tukey HSD *post-hoc* tests, to determine which treatment differed in trait means from the others.

**Table 1:** Results of ANOVA or ANCOVA tests (type II) performed based on linear mixed-effects models with clone genotype as a random factor, and the treatment as fixed factor. These test the effects of treatment on speed of clonal propagation, and morphological and performance traits.

|  | **R2 ramets**<br>(Daughters) |  |  | **R3 ramets**<br>(Granddaughters) |  |  |
| --- | --- | --- | --- | --- | --- | --- |
|  | df | F | P | df | F | P |
| ***Growth*** |  |  |  |  |  |  |
| Speed of clonal propagation | 3 | 81.09 | *** | 3 | 18.55 | *** |
| ***Morphology*** |  |  |  |  |  |  |
| Maximum leaf length | 3 | 32.33 | *** | 3 | 17.21 | *** |
| Maximum root length | 3 | 8.78 | *** | 3 | 5.80 | ** |
| Internode length | 3<br>(1) | 1.50<br>(15.58) | ns<br>(***) | 3<br>(1) | 0.65<br>(14.73) | ns<br>(***) |
| **Performance** |  |  |  |  |  |  |
| Ramet dry mass | 3 | 21.568 | *** | 3 | 24.616 | *** |

Notes: Results into brackets indicate the values obtained for the covariate (ramet dry mass). df = degrees of freedom. F = F-value. P = *p-value*. ns = not significant; \**P* < 0.05; \*\**P* < 0.01; \*\*\**P* < 0.001.

#### Metabolomics data

During our metabolomics analyses on R2 and R3 ramets, we performed both positive and negative ionisation to obtain a wider range of identified features, as both polarities can reveal different metabolites, e.g. negative ionisation is more suitable for acidic compounds (Kim *et al*., 2016). However, many ions detected can also overlap between both ionisations, so to avoid redundancy, we performed all statistical analyses on both datasets independently.

Before all analyses, we applied a cube-root transformation and Pareto scaling for each ionisation dataset. Then, to identify the most important features, we first used a sparse Partial Least Squares (sPLS) approach using the R package *mixOmics* (version 6.24.0) (Rohart *et al*., 2017). To treat both datasets equally, a canonical model was applied. We kept two principal components in our model as only one was optimal, but it takes two to run the model (Supplementary Fig. 1A). Then, the optimal number of features for each component (obtained by using LASSO penalisation) were 40 and 35 for negative ionisation and 45 and 50 for positive ionisation for component 1 and 2 respectively. To test the stability of these selected features, we performed repeated cross-validation (five folds). Beyond this number of selected features, the stability decreased strongly (Supplementary Figs. 1C and 1D), i.e., the quality of the model was not improved by the addition of more features.

To reduce metabolomics data and visualise sample projection, we performed Principal Component Analyses (PCAs) using *FactoMineR* 2.11, and *factoextra* 1.0.7 packages (Lê *et al*., 2008; Kassambara & Mundt, 2016). In addition to PCAs, we also performed Between-Classes Principal Component Analysis (BCA-PCA) using *ade4* 1.7-23 package (Dray & Dufour, 2007) to assess the variance induced by the treatment, quantified by between-class variance (ratio of the amount of variance explained by the sample grouping and the total variance among samples). Between-class variance was tested against simulated values using random permutation test (9999 permutations of the rows of the metabolic feature table) to determine the significant effect of the treatment. Heatmaps were also built to visualise the mean of the top 20 differentially accumulated features in intergenerational metabolic imprint using *gplots* 3.2.0 package (Warnes *et al*., 2005).

All analyses of trait and metabolomics data were performed in R 4.4.2 and RStudio 2024.12.0+467. In addition, volcano plots were done online on BioStatFlow v.2.9.6 © INRAE 2023 to highlight differentially accumulated features (DAFs) between two treatments using *t*-test with False Discovery Rate (FDR) adjustment (fold threshold > 1.5 and *p*-value□<□0.05) and recoloured with Inkscape software.

## Results

### Heat stress reprogramed cellular metabolism

To investigate the metabolome reprogramming resulting from heat stress in our aquatic plant model, *L. australis*, we compared metabolic profiles of R3 ramets in the control (NS), and after a single (NS-S treatment) or repeated episodes of heat stress (S treatment).

First, metabolic profiles of R3 ramets growing at 23 °C (NS-S and S treatments) differed in comparison to those growing at 13 °C (NS treatment) (Figs. 2A and 2B). The two first axes explained 49.2% and 46.7% of total inertia of the PCA, for negative and positive ionisation respectively, and permutation tests confirmed that the treatment had a significant effect on R3 metabolic profiles (negative ionisation: between-class variance□=□0.427, *simulated-P* =□0.001; positive ionisation: between-class variance = 0.394; *simulated-P* =□0.001), indicating metabolome reprogramming after heat stress. To gain further insights into this response, we identified DAFs between R3 ramets of NS and NS-S treatments (Figs. 2C, 2D, 2E, 2F, Supplementary Data 3 and 4). In negative and positive ionisation respectively, 513 (343 annotated) and 1290 (592 annotated) features were differentially accumulated in NS-S treatment while 1148 (906 annotated) and 1538 (756 annotated) were differentially accumulated in NS treatment (Figs. 2C, 2D, Supplementary Data 3 and 4). Regardless of ion modes, most of the features selected belonged to phenylpropanoids and polyketides, lipids and lipid-like molecules, organoheterocyclic compounds, and organic oxygen compounds superclasses (Figs. 2E, 2F, Supplementary Data 3 and 4).

**Figure 2:**
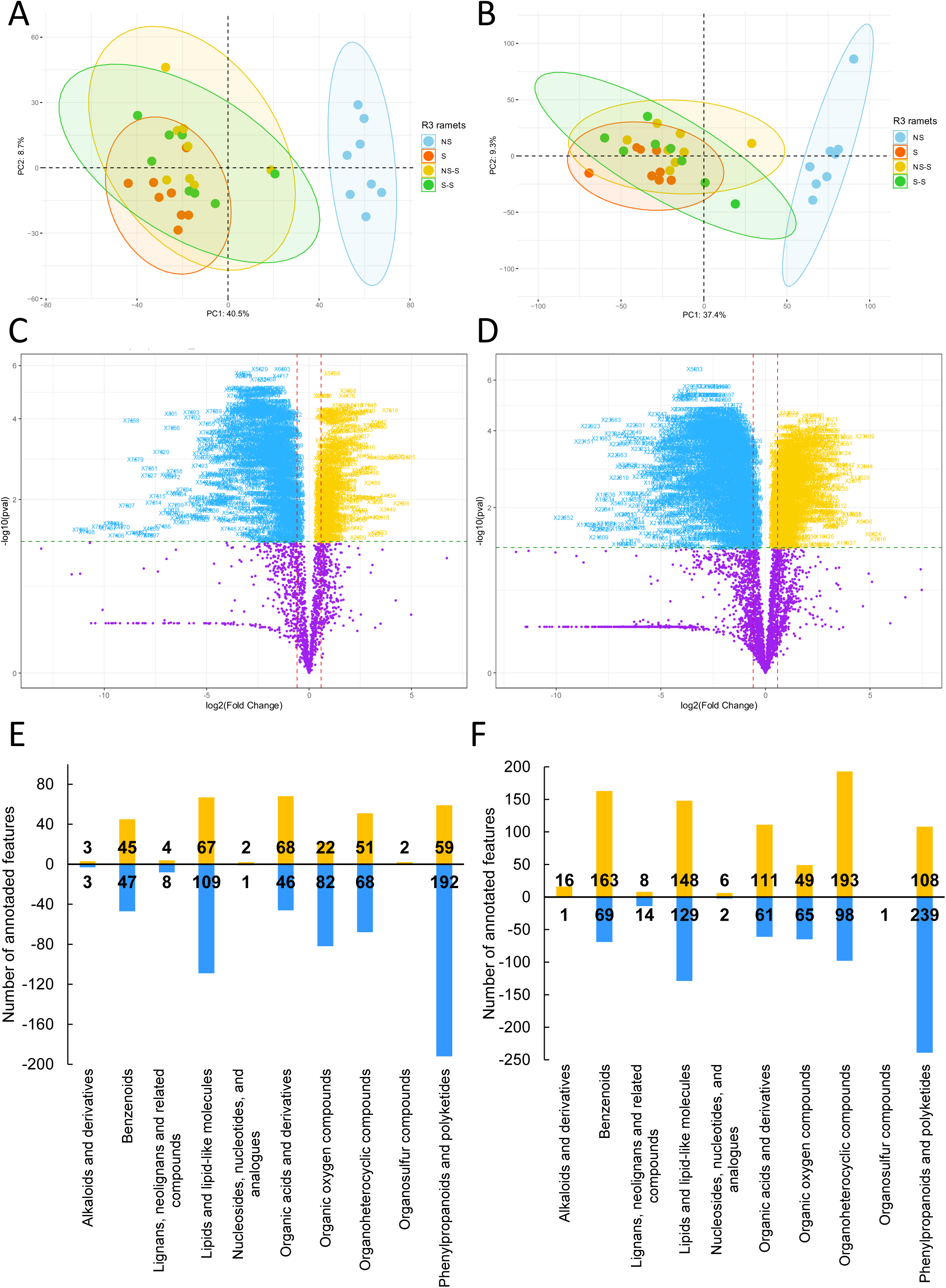
Principal Component Analysis (PCA) score plots of relative features content for granddaughter ramets (R3) coloured by treatment from negative (A) and positive (B) ions mode datasets. Volcano plots of differentially accumulated features (*t*-test with FDR adjustment, fold threshold > 1.5, *p*-value□<□0.05) identified between NS-S and NS treatments (in yellow and blue respectively) in granddaughter ramets for negative (C) and positive (D) ions mode. Number of annotated features in each superclasses that increase (yellow bars) and decrease (blue bars) in NS-S compared to NS treatment for negative (E) and positive (F) ions mode.

Second, we explored the long-term effects of repeated episodes of heat stress and identified DAFs between NS and S treatments in R3 ramets (Supplementary Figs. 2A, 2B, 2C and 2D). In negative and positive ionisation respectively, 550 (369 annotated) and 1384 (972 annotated) features were differentially accumulated in S while 1168 (590 annotated) and 1718 (847 annotated) were differentially accumulated in NS (Supplementary Figs. 2A, 2B, Supplementary Data 5 and 6). Most of the features annotated belonged to the same superclasses as obtained previously, i.e., phenylpropanoids and polyketides compounds or lipids and lipid-like molecules (Figs. 2E, 2F, Supplementary Figs. 2C and 2D). More specifically, over 80% of DAFs between NS and S were similar in both ionisation modes (Supplementary Figs. 2E and 2F).

### Evidence of an intermediate phenotype in daughters of stress-exposed mothers

To determine how the metabolome was modulated by heat stress through the different clonal generations (R2 and R3 ramets), we used a sparse Partial Least Squares (sPLS) approach combining results of both negative and positive ionisations from our untargeted metabolomics analyses (Fig. 3 and Supplementary Fig. 1). For both score plots, we observed a clear separation between ramets living at 13 °C (NS treatment, negative side) and ramets living at 23 °C (S treatment, positive side) along the first axis (Figs. 3A and 3B). Interestingly, we also evidenced an intermediate profile of R2 ramets in S-S compared to R2 ramets in NS and S treatments (Figs. 3A and 3B).

**Figure 3:**
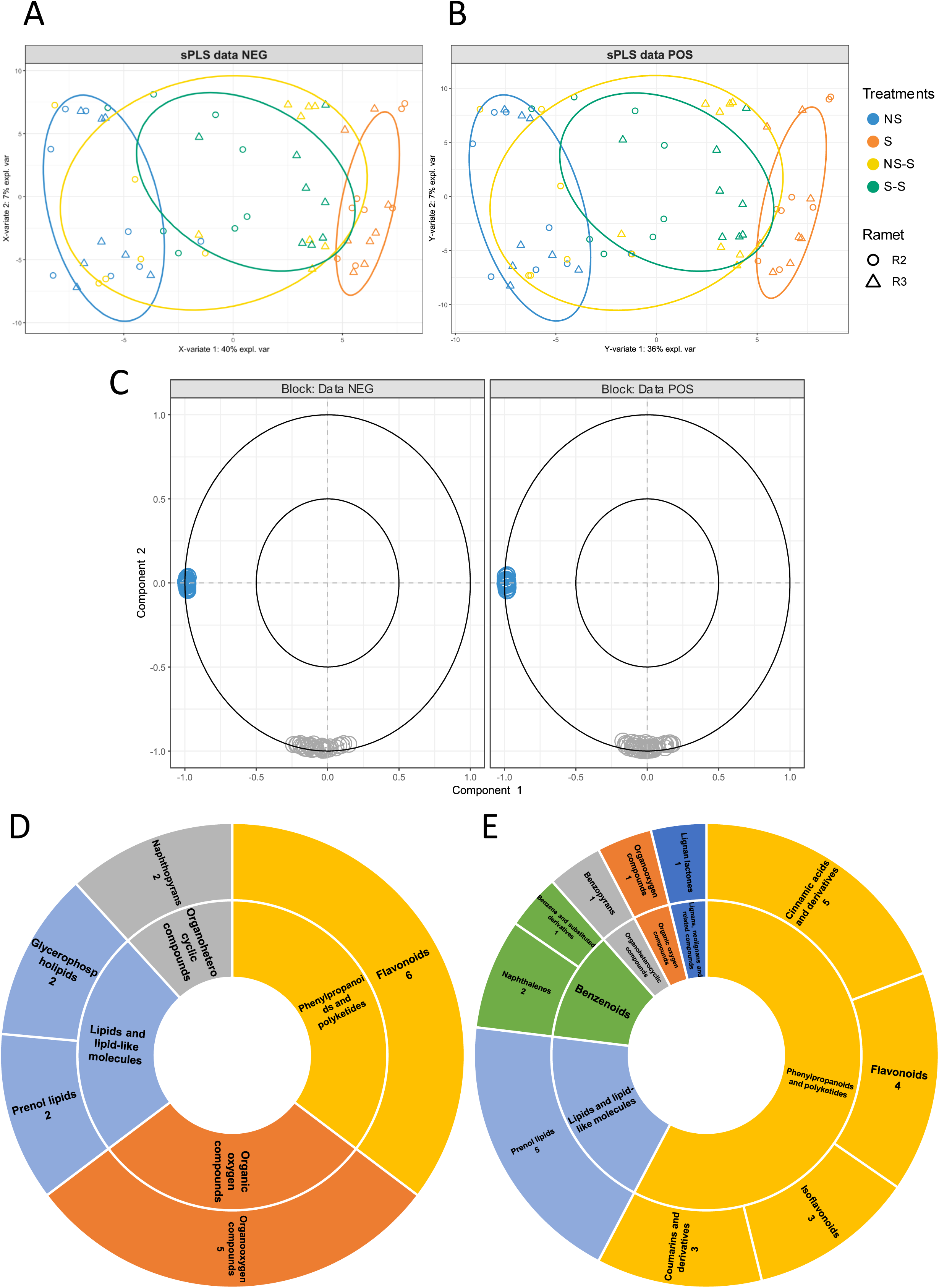
Sample plots for sparse Partial Least Squares (sPLS) analysis performed on negative ionisation data (A) and positive ionisation data (B). Correlation circle plots from sPLS model for each metabolomics data displaying conserved features (C). Sunburst charts of the 22 features annotated of 50 (D) and of the 24 features of 40 (E) (from negative and positive datasets respectively) extracted from component 1 in sPLS models of the corelation circle plots. Class and superclass of features annotated are provided on the inner and outer ring respectively.

Features (40 and 45 for negative and positive ion mode respectively) displayed on the negative side of the first axis corresponded to metabolites differentially accumulated in all ramets stayed at 13 °C except for R2 in S-S treatment (Fig. 3C). Note that no feature selected in our sPLS model was on the positive side of the first axis. We were able to annotate 17 and 26 features out of 40 and 45 for negative and positive ionisation data respectively (Figs. 3D, 3E and Supplementary Data 7 and 8), except for caffeic acid where MS2 information matched (Supplementary Data 8). Most of annotated features belonged to phenylpropanoids and polyketides class, with 10 features belonging to flavonoids superclass, and lipids and lipid-like molecules class (Figs. 3D and 3E).

### Evidence of intergenerational metabolic plasticity

To evidence an intergenerational response resulting from heat stress endured by mother ramet R1, we studied metabolic profiles in R2 ramets (Fig. 4). Metabolic profiles of R2 ramets growing at 13 °C (NS, NS-S and S-S treatments) were significantly different compared to those growing at 23 °C (Figs. 4A and 4B). The two first axes explained 53.9% and 51.8% of total inertia of the PCA, for negative and positive ionisation respectively, and permutation tests indicated that the treatment had a significant effect on metabolic profile (negative ionisation: between-class variance□=□0.482, *simulated-P* =□0.001; positive ionisation: between-class variance = 0.455; *simulated-P* =□0.001). In addition, metabolic profile of S-S R2 ramets also differed from NS and NS-S (Figs. 4A and 4B). As R2 ramets from NS and NS-S experienced the same water temperature from the beginning of the experiment (Fig. 1), we compared and identified DAFs between S-S and NS+NS-S treatments (Figs. 4C and 4D). We identified 648 (316 annotated) and 988 (501 annotated) DAFs, where 128 (69 annotated) and 252 (125 annotated) were accumulated under S-S treatment in negative and positive ionisation respectively (Figs. 4C, 4D, Supplementary Data 9 and 10). Note that we had more features accumulated in NS+NS-S than in S-S treatments (Figs. 4C and 4D). Several superclasses of DAFs were identified, e.g., organic acids and derivatives, phenylpropanoids and polyketides, organoheterocyclic compounds, lipids and lipid-like molecules, or benzenoids, but in each superclasses, most of the features belonged to few different specific classes (Figs. 4E, 4F, Supplementary Data 9 and 10). In negative ionisation, 17 out of the 20 features belonged to carboxylic acids and derivatives class (Fig. 4E) while in positive ionisation, 14 out of the 21 features belonged to prenol lipids class (Fig. 4F). We also identified several flavonoids in both ionisation analyses (Supplementary Data 9 and 10). Finally, especially in negative ionisation, we also identified some features annotated as amino acids or amino acid derivates, like aspartic acid, isoleucine, glutamic acid, glutamine and tryptophan (Supplementary Data 9).

**Figure 4:**
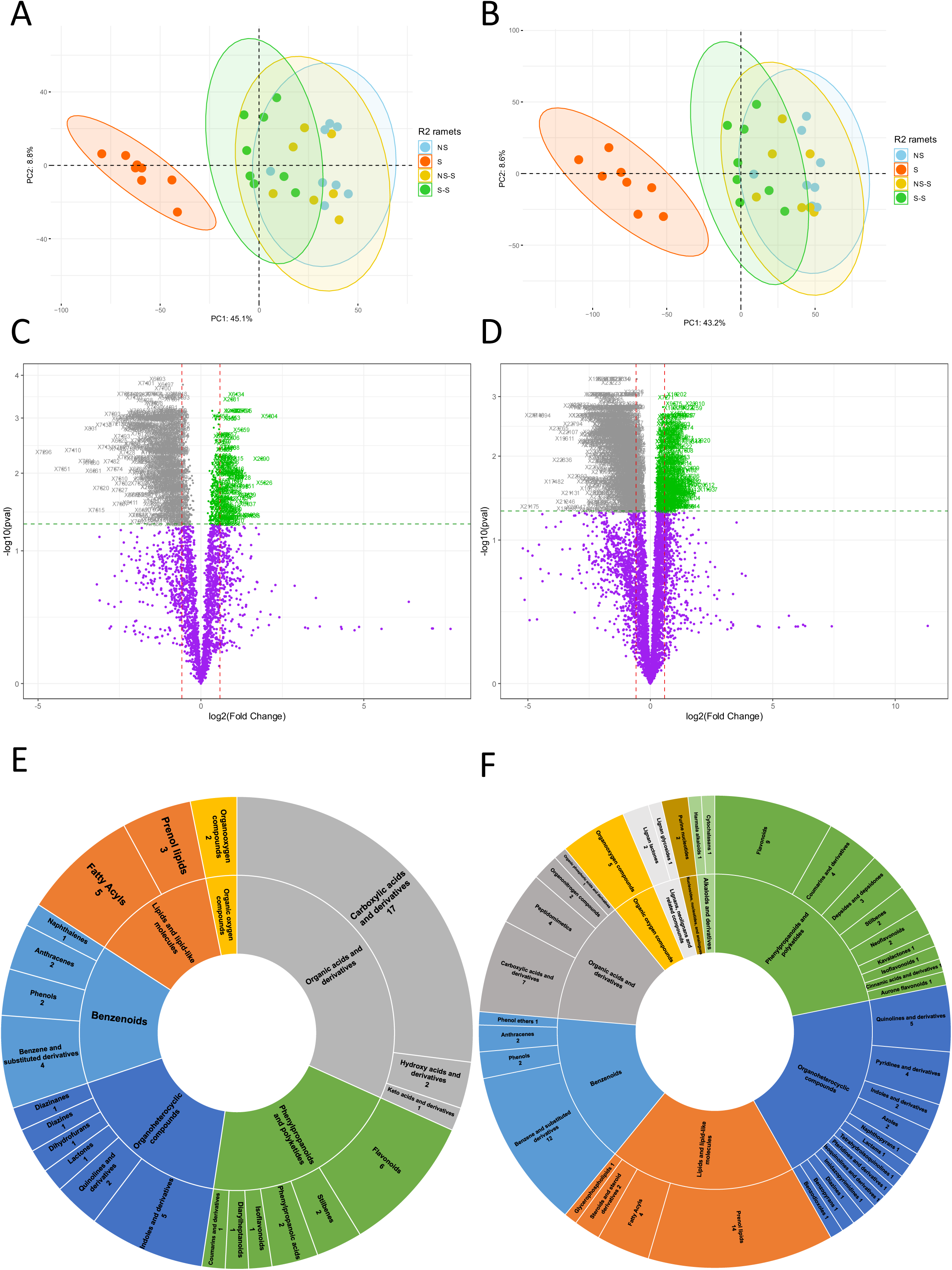
Principal Component Analysis (PCA) score plots of relative features content for daughter ramets (R2) coloured by treatment from negative (A) and positive (B) ions mode datasets. Volcano plots of differentially accumulated features (*t*-test with FDR adjustment, fold threshold > 1.5, *p*-value□<□0.05) identified between S-S and NS+NS-S treatments (in green and grey respectively) in daughter ramets for negative (C) and positive (D) ions mode. Sunburst charts of number of annotated features that increase in S-S compared to NS+NS-S in daughter ramets for negative (E) and positive (F) ions mode. Class and superclass of features annotated are provided on the inner and outer ring respectively.

To investigate how plant memory affects the metabolic response of granddaughter R3 ramets during a second episode of heat stress, we characterised their metabolic profiles. However, regarding metabolic profiles of R3 ramets in S-S and NS-S treatments, we could not find any DAFs in both ionisation analyses (Supplementary Fig. 3).

### Metabolic imprint induced by heat stress was highly conserved in daughter ramets

To further understand the induction of the potential metabolic markers identified during the recovery period in S-S treatment in daughter ramets (R2), we compared them to features induced during a first heat stress (Figs. 5A and 5B), previously identified in NS-S *vs.* NS and S *vs.* NS treatments in R3 ramets (Fig. 2 and Supplementary Fig. 2). Many of them were also induced during a first heat stress (Fig. 5), indicating that these common features remained accumulated after the heat stress during the recovery period in the next clonal generation (R2, daughter ramets). The other features constituting this metabolic imprint in daughter ramets, were then S-S-specific, i.e. their accumulation was found only in the next clonal generation (Figs. 5A and 5B). This was the case for four unknown features (id_7639, id_3352, id_5659 and id_11687) in the top20 (the 20 highest-ranking features with fold change >1.5), where a higher accumulation was identified only in S-S treatment in daughter ramets R2 (Figs. 5C and 5D).

**Figure 5:**
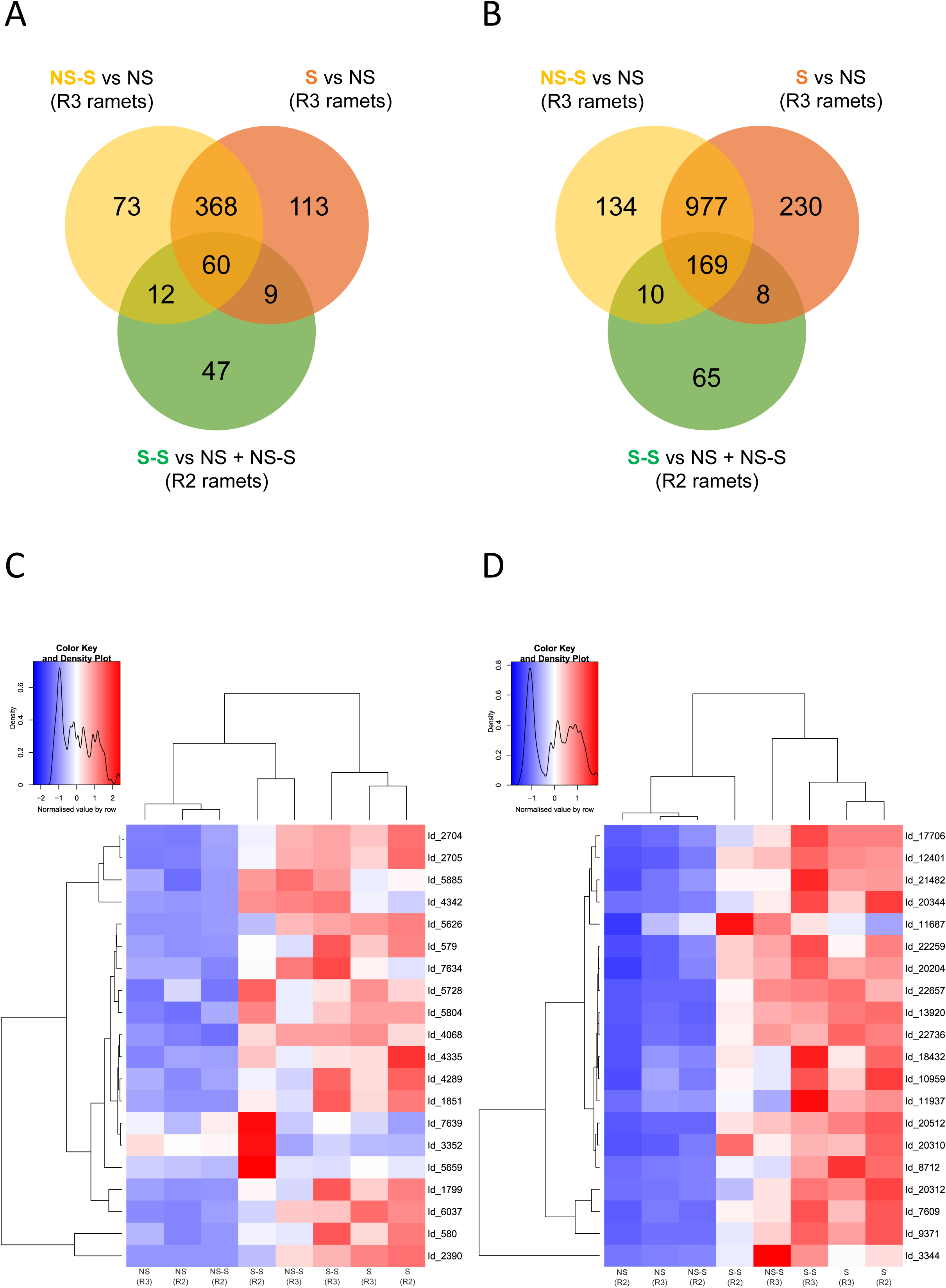
Venn diagrams of the number of differentially accumulated features between NS and NS-S (granddaughter ramets), between NS and S (granddaughter ramets), and between NS+NS-S and S-S (daughter ramets), for negative (A) and positive (B) ionisation modes. Heatmaps of the relative content of top20 features (the 20 highest fold threshold) that accumulated in S-S compared to NS+NS-S in all treatments and ramets studied for negative (C) and positive (D) ions mode.

### Intergenerational metabolic responses led to modifications in daughter growth, morphology, and performances

To investigate the effect of mother environment on daughter and granddaughter development, we measured the speed of clonal propagation, and the morphological and performance traits on R2 and R3 ramets (Fig. 6 and Table 1).

**Figure 6:**
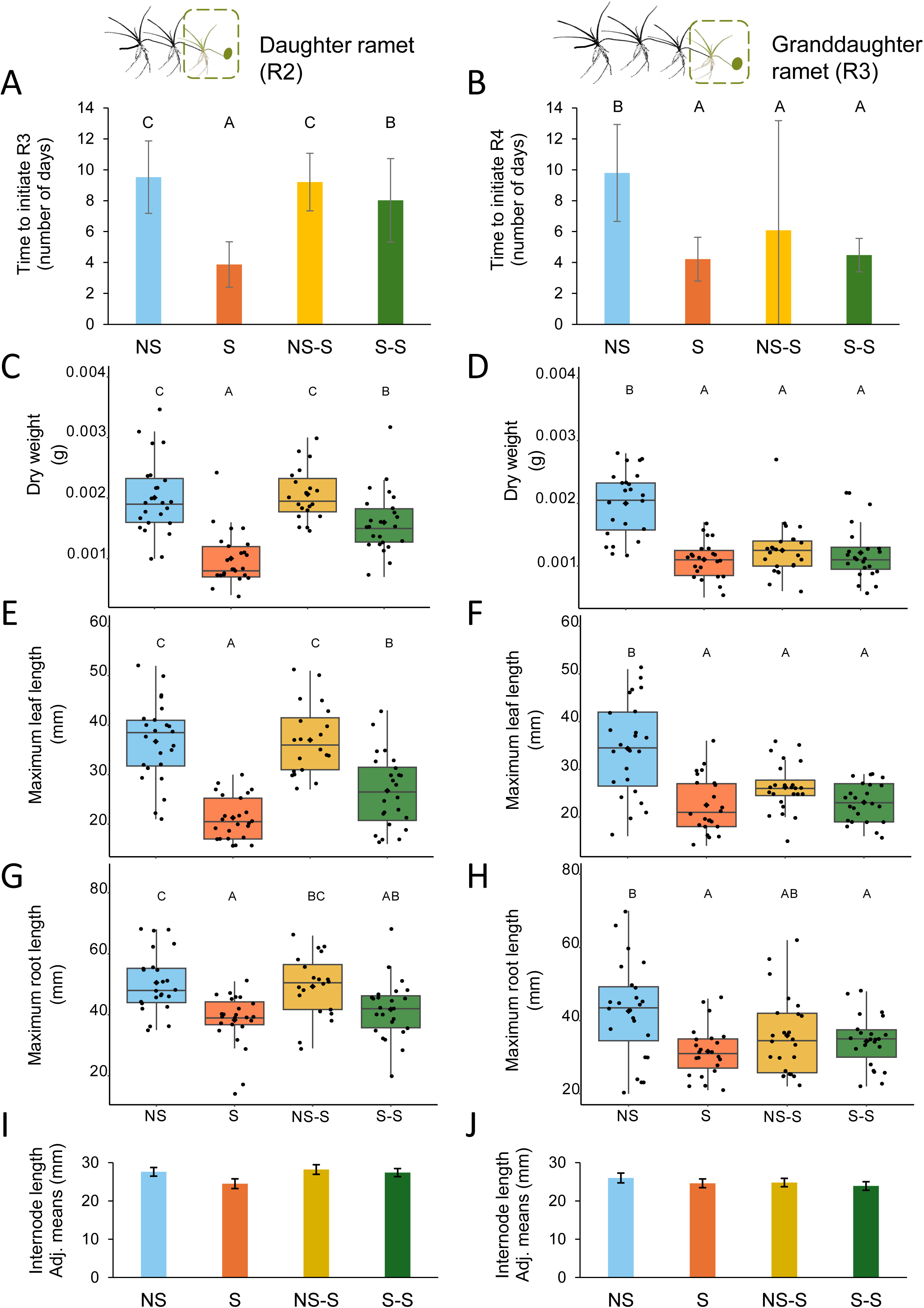
Measured traits to characterise growth, morphology and ramet performance in daughter and granddaughter ramets (R2 and R3). Means and standard deviation for the number of days for daughter ramets to produce a new clonal ramet (R3) in the four treatments applied (A). Means and standard deviation for the number of days for granddaughter ramets to produce a new clonal ramet (R4) in the four treatments applied (B). Boxplots displaying dry weight, maximum leaf length and maximum root length of R2 (C, E and G) and R3 (D, F and H) ramets. Adjusted means and standard error of internode length of R2 (I) and R3 (J) ramets using dry weight as a covariate effect (least square means predicted from the models). Letters are based on Adj. means and indicate significant differences between treatments using pairwise comparison post-hoc tests with Tukey method *p-value* adjustment.

Comparing R3 growth and performances between S-S and NS-S treatments, we did not evidence any significant differences (Figs. 6B, 6D, 6F, 6H and 6J).

In R2 however, morphological traits (except for internode length, Fig. 6I) and speed of clonal propagation were significantly influenced by the treatment (Table 1). Under NS and S stable conditions, we observed that R2 maximum leaf length and maximum root length were longer (Figs. 6E and G) and that R2 ramet initiated R3 faster (Fig. 6A) at 23 °C than at 13 °C. In S-S treatment, R2 ramets displayed an intermediate phenotype compared to NS, S and NS-S treatments (Figs. 6A, 6E and 6G). Regarding performances, at 23 °C, ramet dry weight was lower in S compared to NS, and intermediate under S-S treatment (Fig. 6C, Table 1).

## Discussion

### An increase in water temperature implied huge shift in metabolic profile within few days

In line with our first hypothesis, our results demonstrated that heat stress strongly affected *L. australis* metabolome, including both primary and secondary metabolic pathways. Surprisingly, we observed an overall decrease of features content during the heat stress, including lipids, organic acids and phenylpropanoid compounds. This result appeared counter-intuitive as we rather expected an increase in feature content at 23 °C, based on the literature demonstrating that heat stress usually promotes metabolite accumulation (Guy *et al*., 2008; Du *et al*., 2011). It could be explained by the higher growth rate of clones at 23 °C than at 13 °C. As 23 °C ramets were produced more rapidly, they did not accumulate metabolites as much as ramets at 13 °C.

Despite the wide diversity of features found, several amino acids (e.g. tryptophan, isoleucine, lysine, pyroglutamate or methionine) and secondary metabolites (e.g., flavonoids) were annotated. For example, tryptophan is essential to protein biosynthesis and participates in regulating plant growth and defence through auxin accumulation (Liu *et al*., 2022). Modifications in amino acids content may thus have led to the higher speed of clonal propagation observed in R3 ramets for NS-S and S treatments. In addition, flavonoids are known for their multiple cellular functions, especially for their antioxidant activity (Agati *et al*., 2012), but also for their role in signalisation in various biochemical pathways (Taylor & Grotewold, 2005). So, the lower flavonoids accumulation may decrease the defence status of ramets living at 23 °C, as they normally increase during thermal stresses (Ren *et al*., 2026).

Comparison of metabolic responses to short- (R3 in NS-S) and long-term (R3 in S) heat stress showed very little differences. Most of the metabolic changes were similar, suggesting that only one clonal generation is needed for *L. australis* to respond to a warmer aquatic environment. Heat stress triggers molecular responses within several seconds to minutes (Kollist *et al*., 2019). However, under moderate heat stress over long term, these responses decrease as plants acclimate to a new physiological balance (Tang *et al*., 2016). Thus, because metabolic responses were similar between NS-S and S conditions in R3 ramets and most of secondary metabolites decreased in comparison to NS condition, we proposed an alternative hypothesis that an increase in water temperature by 10 °C (from 13 to 23 °C) was actually perceived as moderate heat stress by *L. australis*, allowing it to respond within a few days. Furthermore, *L. australis* is distributed across several latitudes and lives under warmer climates than in the Iles Kerguelen, e.g. south Australia and east coast of south America (MNHN & OFB [Ed], 2003-2025), suggesting that this species is actually eurytherm. The effect of climate change is already visible on the Iles Kerguelen (Leihy *et al*., 2018) with higher annual temperatures and fewer frost days (Frenot *et al*., 2006). We therefore expect *L. australis* to be able to survive and persist despite the increase of heat stress events in the future, both at the freshwater pond scale and across the whole archipelago.

### Metabolome profiling of daughter ramets demonstrated an intergenerational metabolic plasticity

Understanding the importance of inter- and transgenerational plasticity in response to changing environment is crucial as it may support species ability to persist under the novel climatic conditions (higher and more fluctuating temperatures) resulting from climate change. More specifically, insights into inter- and transgenerational metabolic memory are needed and complementary to genomic and proteomic studies to characterise metabolic pathways involved in plant memory.

Our work focused notably on the identification of an intergenerational plasticity, and in line with our second hypothesis, our results highlighted that maternal environment influenced daughter ramet metabolism and revealed an intergenerational metabolic plasticity. In fact, we observed an intermediate profile in our metabolomics analyses, with a separation between S-S, NS/NS-S and S treatments in R2 ramets. These differences of relative feature concentrations in daughter ramets corresponded to an intergenerational imprint of the mother environment in S-S treatment, during the recovery period. Among these features, we could find several organic acids, and more specifically some amino acids, like isoleucine or glutamic acid, confirming typical signatures of past stress effects found in previous studies (Mandal *et al*., 2012; Benina *et al*., 2013; An *et al*., 2013; Wedeking *et al*., 2018; Auler *et al*., 2021). Although we focus on DAFs in S-S treatment, we observed a global reduction of relative feature contents compared to NS/NS-S treatments. This outcome was not surprising as we also observed a global feature reduction during heat stress in mothers. In addition, a recent study observed similar results identifying a decrease in Shannon diversity (used to assess metabolome diversity) in *Festuca rubra* clonal ramets by comparing offspring from recurrent drought-stressed parents with offspring from stress-free parents (Bhatt *et al*., 2026). Even if there probably are common mechanisms of metabolic memory among plants, like an amino acid accumulation (Zeier, 2013), each species also has its own specific stress and post-stress responses (Mandal *et al*., 2012; Benina *et al*., 2013). We then highlighted that many other metabolites were part of the metabolic imprint. These other features were annotated into several classes and superclasses, including carboxylic acids, prenol lipids like terpenes, fatty acids, or phenylpropanoids like flavonoids, highlighting the wide range of metabolites potentially involved in metabolic memory. By comparing the accumulation of these different features with those accumulated during heat stress, we demonstrated that few compounds accumulated specifically in daughter ramets. This was the case for two features with MS2 confirmation, tryptophan and kaempferol-3-glucuronide. Conversely, and interestingly, most other features from the metabolic imprint were also found to be accumulated during mother heat stress, like pyroglutamate and isoleucine, meaning that they stayed in high concentrations in daughters during the recovery period. These findings are partially consistent with another study suggesting different metabolic responses in *Chlamydomonas reinhardtii* during the recovery period following a long term heat stress (Hemme *et al*., 2014). More specifically, Hemme *et al*., 2014 mostly evidenced that compounds from Krebs cycle, like succinate, malate, or citrate, accumulated in higher concentrations during the recovery period than before or during the heat stress, while other compounds like isoleucine and other amino acids accumulated during heat stress and stayed in high concentration during the recovery period (Hemme *et al*., 2014). Thus, in addition to having identified diverse metabolite families potentially involved in metabolic memory, we also demonstrated that some compounds can be accumulated during heat stress and stayed in high concentration in daughters, while a few were specifically accumulated during the recovery period.

Studies of transgenerational plasticity in clonal plant remains rare, in particular those dealing with a second stress event (Zhang *et al*., 2023; Jin *et al*., 2023), and to our knowledge, there are no studies characterising the metabolome at the transgenerational level. At last, our work also sought to highlight transgenerational plasticity, and how it affects granddaughter metabolome in response to a second heat stress. However, and contrary to our third hypothesis, we did not evidence transgenerational metabolic plasticity.

### Growth strategy shifted during heat stress and persisted in the next clonal generation

Our results support that heat stress exposure in mothers affects their growth strategy, influenced the daughter’s performance and morphology, but had no effect on granddaughters exposed to heat stress, partially validating hypothesis 4.

More specifically, under heat stress, mothers showed an overall decrease in ramet performance, i.e. lower dry weight, shorter maximum length of leaves and roots and faster initiation of new clonal ramet. This outcome suggested that mother ramets first allocated resources to a more rapid production of new daughters. Note that similar findings have been previously observed for this species (Douce *et al*., 2024), although numerous studies examining temperature influence on plant morphology and performance, highlighted contrasting responses depending on the range of temperatures and on the thermal optimum of the studied plant species (Pilon & Santamaría, 2002; Riis *et al*., 2012; Chalanika De Silva & Asaeda, 2017; Beca-Carretero *et al*., 2018; Wittyngham *et al*., 2019; Lauridsen *et al*., 2020; Billah *et al*., 2025). However, despite a decrease of ramet performance, we thought that this is partly compensated at the clone level (Douce *et al*., 2024). Furthermore, in our study, internode length remained the same at 13 °C and 23 °C, characterising an active process of internode elongation at 23 °C despite lower ramet performance. In clonal plants, a decrease or an increase of internode length gives us a clue about plant foraging behaviour (Louâpre *et al*., 2012). We thus suggested that plant growth strategy switched to a more efficiently colonise space, by reducing ramet performance, maintaining mass allocation to internode elongation, and speeding up ramet initiation.

We explored whether these plastic responses to heat stress observed in the mothers were transmitted to the progeny through intergenerational plasticity. Intergenerational plasticity in response to changing environments can drive the development of new clonal generations (González *et al*., 2017; Münzbergová & Hadincová, 2017; Portela *et al*., 2020; Quan *et al*., 2022), subsequently influencing the species’ capacity to adapt to novel climatic conditions (Münzbergová & Hadincová, 2017). We observed that heat stress in maternal generation decreased daughter growth and performance, and increased the speed of clonal propagation but without reaching the values found in stressed mothers. This has already been partially observed for particular levels of drought stresses in the clonal plant *Trifolium repens* (González *et al*., 2016). In other words, heat stress experienced by mothers influenced daughter development, resulting in an intermediate foraging strategy between stressed and stressed daughters (from non-stressed mothers). This may reduce the individual costs if another heat stress occurs. From an ecological point of view, this reinforced the idea that ramets of *L. australis* species will likely survive and maintain their presence within the macrophyte communities at the Iles Kerguelen.

### Transgenerational effects of heat stress did not provide any adaptative advantage

Contrary to the intergenerational response that we observed during the recovery period, heat stress on mothers did not provide any adaptative (or either maladaptive) advantage to granddaughter exposed to a second heat stress. Similar results could be found in different clonal plants in response to herbivory or different temperature conditions (González *et al*., 2017; Münzbergová & Hadincová, 2017). Nevertheless, it has been found that transgenerational responses to plant competition was maladaptive in a perennial floating clonal plant, *Spirodela polyrhiza* (L.) Schleid (Jin *et al*., 2023). Thus, our findings suggested that, in *L. australis*, transgenerational effects do not persist beyond the first clonal generation, despite that these could persist at least for two clonal generations in other clonal plant species (Zhang *et al*., 2023; Jin *et al*., 2023). This suggestion was supported by the absence of DAFs identified in granddaughters, showing that even at the metabolome level, we were not able to find any metabolic imprint from this past heat stress. To explain this result, we suggested two non-mutually exclusive hypotheses. First, a recovery period lasting for an entire clonal generation was enough to fully reset, or at least dilute, all transgenerational effects. That type of short-term memory could allow daughters to rapidly respond to heat stress, without maintaining a lower ramet performance across more than one generation during suitable conditions. Second, inter- and transgenerational memory in clonal plants, is directly link to their distance of physiological integration within the clone (Liu *et al*., 2016; Wang *et al*., 2021). It is then possible that this distance is short in *L. australis*, and limited to one connected ramet. Thus, the memory mechanisms and signals from heat stressed mothers might not have been inherited by the granddaughters, but by the daughters only.

Despite the importance of inter- and transgenerational plasticity for clonal plants to cope with fluctuating environment, highly variable and non-predictable environmental conditions, especially at short-time scales, should favour non-plastic strategies (in our study, plastic response to heat stress persisted for one generation only, i.e. low plasticity at the clone level) as plasticity could lead to environment-phenotypes mismatch (Magyar *et al*., 2007). Temperatures in freshwater ponds on the Iles Kerguelen can be highly variable even at a daily scale (Douce *et al*., 2023), suggesting that *L. australis* may have an adaptive advantage to not keep a stress memory over several clonal generations.

To conclude, our findings bring new insights into the responses to climate change of aquatic plants living in polar regions or cold ecosystems. In particular, the maintenance (or the loss) of transgenerational plasticity in these plants may have strong implications for their fate in a context of more frequent abiotic stresses.

## Acknowledgements

The authors thank Jeremy Bacon and Lara Konecny for plant sampling and laboratory support.

## Author contribution

AKB and GL designed the experiment; MS and GL cultivated the plants, measured plant traits, prepared samples and extractions; JVF, PP and GL did the untargeted metabolome analysis and processed it with MS-DIAL software (v4.9); GL analysed and performed statistical analysis of the data; GL wrote the manuscript; AKB revised and corrected the manuscript. All authors read and approved the final manuscript.

## Data availability

Raw metabolomics data are available upon request.

Metadata and processed data are given in Supplementary Data 1 and 2.

---

**Supplementary Figure 1:**
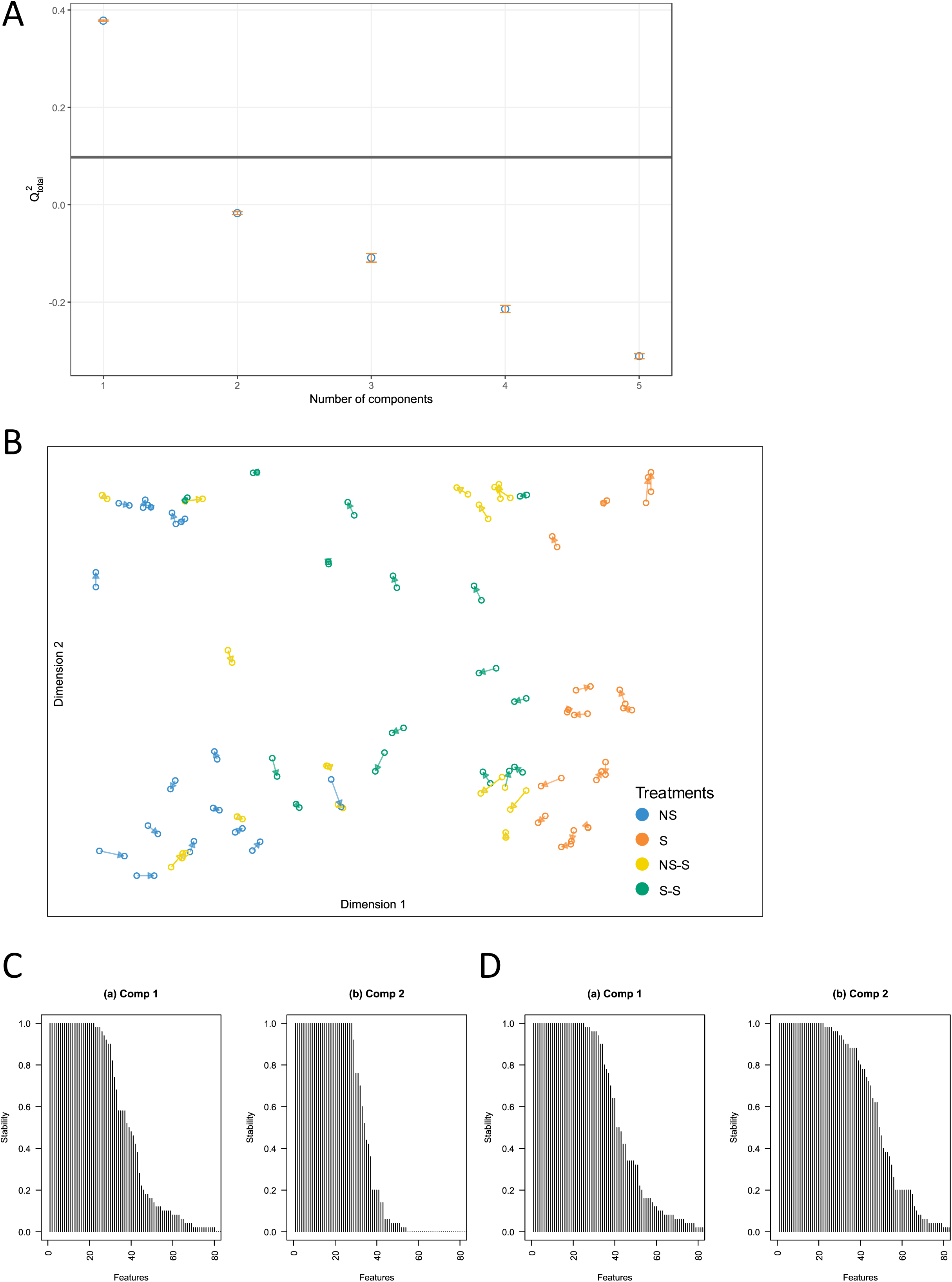
Tuning the number of components in sPLS. For each component, Q² score is shown (repeated cross-validation, 5 × 10−fold CV). Error bars represent value variations across the repeated folds (A). Arrow plot from sPLS model. The start and the tip of the arrow indicate the same sample in the space spanned by the components associated to negative and positive ionisation dataset respectively (B). Stability of feature selection from the sPLS on negative (C) and positive (D) ionisation dataset (repeated cross-validation, 5 × 10−fold CV).

**Supplementary Figure 2:**
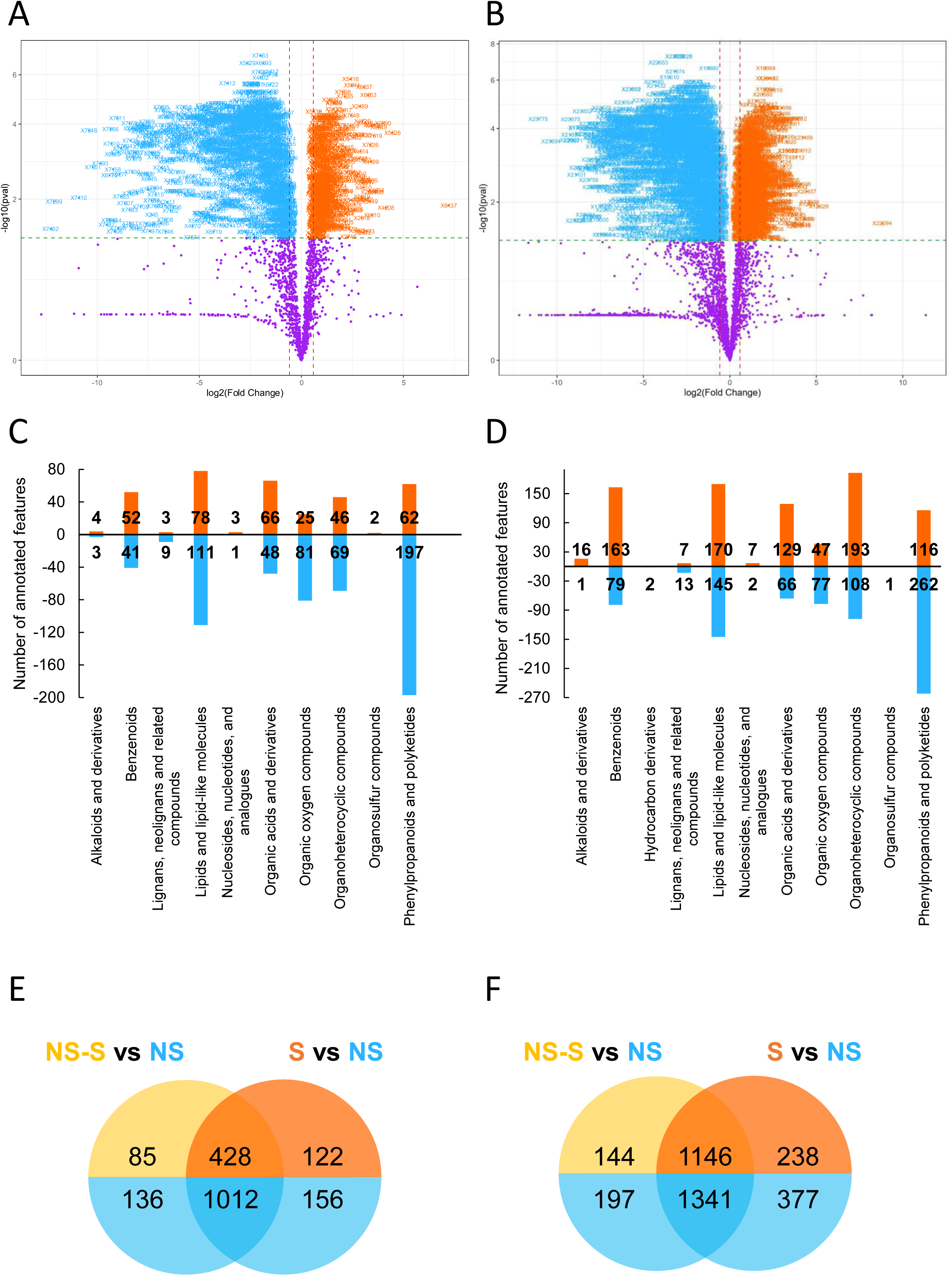
Volcano plots of differentially accumulated features (*t*-test with FDR adjustment, fold threshold > 1.5, *p*-value□<□0.05) identified between S and NS treatments (in orange and blue respectively) in granddaughter ramets (R3) for negative (A) and positive ions mode (B). Number of annotated features in each superclasses that increase (orange bars) and decrease (blue bars) in S compared to NS for negative (C) and positive (D) ions mode. Venn diagrams of the number of differentially accumulated features between NS and NS-S and between NS and S conditions in granddaughter ramets (R3) for negative (E) and positive (F) ionisation modes.

**Supplementary Figure 3:**
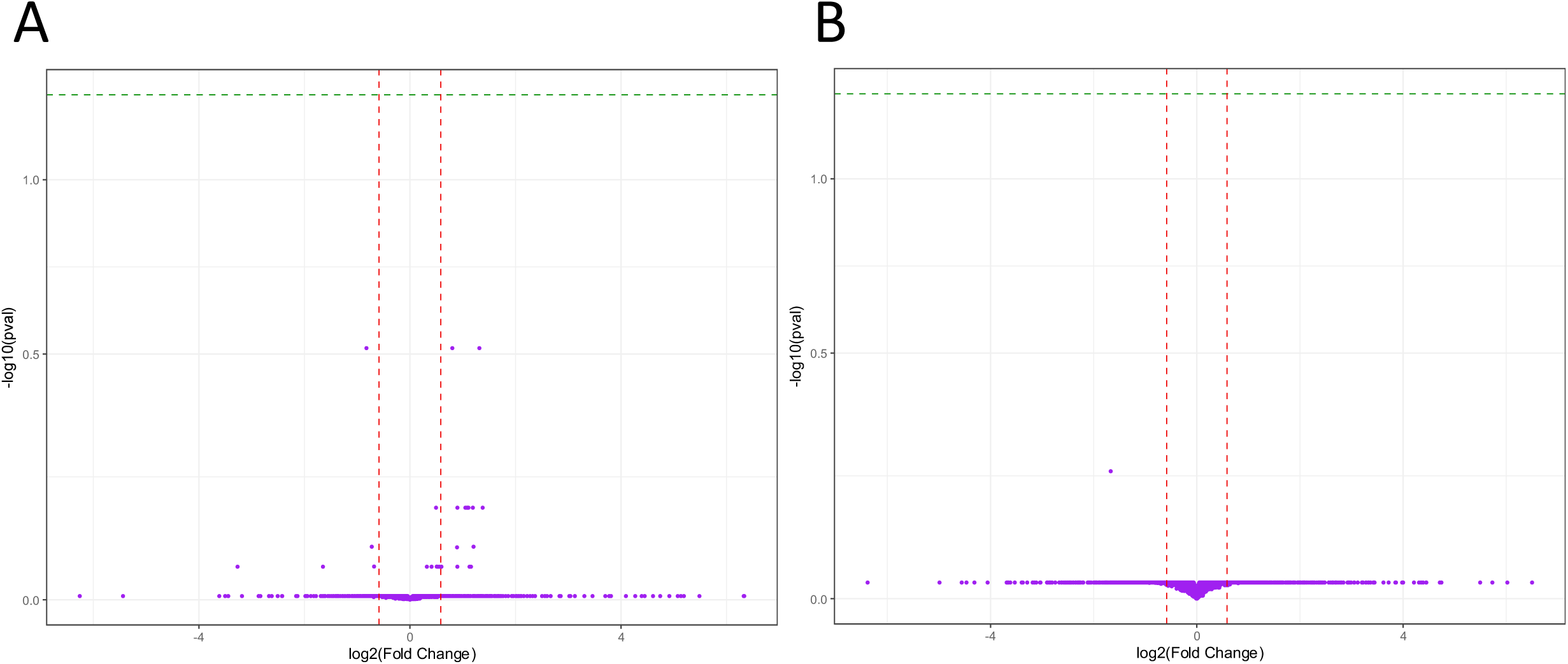
Volcano plots of differentially accumulated features (*t*-test with FDR adjustment, fold threshold > 1.5, *p*-value□<□0.05) identified between S-S and NS-S conditions in granddaughter ramets (R3) for negative (C) and positive (D) ions mode.

Supplementary Data 1: Relative content for the 3267 features extracted from negative untargeted metabolic analysis and their annotated chemical ontology.

Supplementary Data 2: Relative content for the 5942 features extracted from positive untargeted metabolic analysis and their annotated chemical ontology.

Supplementary Data 3: Differentially accumulated features in NS-S treatment compared to NS in granddaughter ramets (R3) (*t*-test with FDR adjustment, fold threshold > 1.5, *p*-value□<□0.05) from negative ionisation dataset and their annotated chemical ontology.

Supplementary Data 4: Differentially accumulated features in NS-S treatment compared to NS in granddaughter ramets (R3) (*t*-test with FDR adjustment, fold threshold > 1.5, *p*-value□<□0.05) from positive ionisation dataset and their annotated chemical ontology.

Supplementary Data 5: Differentially accumulated features in S treatment compared to NS in granddaughter ramets (R3) (*t*-test with FDR adjustment, fold threshold > 1.5, *p*-value□<□0.05) from negative ionisation dataset and their annotated chemical ontology.

Supplementary Data 6: Differentially accumulated features in S treatment compared to NS in granddaughter ramets (R3) (*t*-test with FDR adjustment, fold threshold > 1.5, *p*-value□<□0.05) from positive ionisation dataset and their annotated chemical ontology.

Supplementary Data 7: The list of features extracted from sPLS model for component 1 from negative ion mode analysis and their annotated chemical ontology.

Supplementary Data 8: The list of features extracted from sPLS model for component 1 from positive ion mode analysis and their annotated chemical ontology.

Supplementary Data 9: Differentially accumulated features in S-S treatment compared to NS+NS-S in daughter ramets (R2) (*t*-test with FDR adjustment, fold threshold > 1.5, *p*-value□<□0.05) from negative ionisation dataset and their annotated chemical ontology.

Supplementary Data 10: Differentially accumulated features in S-S treatment compared to NS+NS-S in daughter ramets (R2) (*t*-test with FDR adjustment, fold threshold > 1.5, *p*-value□<□0.05) from positive ionisation dataset and their annotated chemical ontology.

Supplementary File 1: Sample list for negative and positive untargeted metabolomics analyses, indicating raw sample names, blanks, QC and samples, the analytical order, name of treatment, the clone genotype, and the biomass (ramet dry weight) for data normalisation.

Supplementary File 2: Parameters used to processed raw data via MS-DIAL software for negative ionisation data acquisition.

Supplementary File 3: Parameters used to processed raw data via MS-DIAL software for positive ionisation data acquisition.

